# Loss of Skeletal Muscle Pyruvate Dehydrogenase Induces Lactic Acidosis and Adaptive Anaplerotic Compensation via Pyruvate-Alanine Cycling and Glutaminolysis

**DOI:** 10.1101/2023.05.10.540277

**Authors:** Keshav Gopal, Abdualrahman Mohammed Abdualkader, Xiaobei Li, Amanda A. Greenwell, Qutuba G. Karwi, Christina Saed, Golam M. Uddin, Ahmed M. Darwesh, K Lockhart Jamieson, Tariq R. Altamimi, Ryekjang Kim, Farah Eaton, John M. Seubert, Gary D. Lopaschuk, John R. Ussher, Rami Al Batran

## Abstract

Pyruvate dehydrogenase (PDH) is the rate-limiting enzyme for glucose oxidation that links glycolysis-derived pyruvate with the TCA cycle. Although skeletal muscle is a significant site for glucose oxidation and is closely linked with metabolic flexibility, the importance of muscle PDH during rest and exercise has yet to be fully elucidated. Here, we demonstrate that mice with muscle-specific deletion of PDH exhibit rapid weight loss and suffer from severe lactic acidosis, ultimately leading to early mortality under low-fat diet provision. Furthermore, loss of muscle PDH induces adaptive anaplerotic compensation by increasing pyruvate-alanine cycling and glutaminolysis. Interestingly, high-fat diet supplementation effectively abolishes the early mortality and rescues the overt metabolic phenotype induced by muscle PDH deficiency. Despite increased reliance on fatty acid oxidation during high-fat diet provision, loss of muscle PDH worsens exercise performance and induces lactic acidosis. These observations illustrate the importance of muscle PDH in maintaining metabolic flexibility and preventing the development of metabolic disorders.

**Highlights:**

- Skeletal Muscle PDH is essential for survival
- Loss of muscle PDH induces lactic acidosis and premature death
- Loss of muscle PDH enhances pyruvate transformations and glutaminolysis
- High-fat diet supplementation abolishes early mortality and overt phenotype induced by muscle PDH loss

## INTRODUCTION

Metabolic flexibility describes the ability of an organism to switch between different fuel sources and metabolic pathways to meet energy needs under various conditions^1^. This concept has been positioned at the forefront of diabetes research for several decades to explain the mechanisms underlying muscle insulin resistance and the metabolic energy switch that occurs in obesity and type 2 diabetes. In 1963 Randle and colleagues postulated the glucose-fatty acid cycle, also known as the Randle cycle, as an attempt to describe the competition between fatty acids and glucose for their oxidation and uptake in muscle and adipose tissue^2^. In this model, they proposed that increased fatty acid oxidation produces acetyl-CoA and NADH, which in turn allosterically inhibits pyruvate dehydrogenase (PDH), the rate-limiting enzyme for glucose oxidation that couples glycolysis to glucose oxidation. They further suggested that the accumulation of citrate resulting from enhanced fatty acid oxidation would inhibit phosphofructokinase leading to decreased glycolysis and increased intracellular glucose 6-phosphate concentrations, thus inhibiting hexokinase and glucose uptake.

Randle’s findings spawned an enormous amount of research over the following four decades to validate this hypothesis. One of the most significant attempts was by Kelley and Mandarino^3^, who conducted a series of influential studies on healthy individuals, demonstrating that muscles primarily rely on fatty acid oxidation in the post-absorptive state and glucose oxidation during the fed state. However, in patients with obesity and type 2 diabetes, muscles rely on fatty acid oxidation even in the fed state, and become unresponsive to insulin action^4-6^. These studies coined the term metabolic inflexibility, a phenomenon that is strongly linked to insulin resistance and metabolic syndrome but remains largely unknown and merits investigation. Nevertheless, with the rising prevalence of obesity worldwide, multiple hypotheses were put together in an effort to pinpoint potential targets that could alter energy substrate preference and reverse metabolic inflexibility and insulin resistance. For example, one model suggested that prolonged muscle exposure to fatty acids leads to incomplete fatty acid β-oxidation and insulin resistance, thereby impairing the muscle’s ability to switch to glucose utilization during the transition from a fasted to a fed state. However, restricting mitochondrial fatty acid uptake can potentially reverse these effects and improve metabolic flexibility^7^.

Although these studies clearly demonstrate that glucose oxidation is a key determinant of metabolic flexibility, the importance of muscle PDH in regulating energy substrate selection based on substrate availability and energy demands has not been fully elucidated. In an attempt to shed light on this question, we examined the impact of skeletal-muscle-specific PDH deficiency on muscle fuel selection at rest and during exercise.

## RESULTS

### Loss of Muscle PDH Induces Lactic Acidosis and Premature Mortality

Because global knockout of PDH is embryonically lethal^8^ and mice with a skeletal muscle- and heart-specific PDH deficiency die within 7 days of weaning^9^, we generated mice with a tamoxifen-inducible skeletal muscle–specific knockout of PDH (PDH^SkM-/-^). Skeletal muscle-specific deletion of PDH was confirmed with western blot analysis, where PDH protein expression was absent from PDH^SkM-/-^ muscles while preserved in the heart (**Figure 1A**). Interestingly, deleting muscle PDH did not affect the expression of genes that regulate the PDH complex, namely PDH kinase (*Pdk*) and PDH phosphatase (*Pdp*) isozymes (**Supplementary Figure 1A**). PDH^SkM-/-^ mice were indistinguishable from HSA^*Cre*^ mice littermates pre-tamoxifen injections and PDH deletion. Notably, ∼1 week post tamoxifen-induced skeletal muscle PDH deletion, PDH^SkM-/-^ mice fed a low fat diet (LFD) became overtly symptomatic, appeared wasted and weak, and ultimately died within 3 weeks post-tamoxifen administration (**Figure 1B**). We noticed that PDH^SkM-/-^ mice exhibited rapid weight loss that was manifested by significant reduction in lean mass and total body length compared to their HSA^*Cre*^ littermates (**Figure 1C-E**). Furthermore, PDH^SkM-/-^ mice when subjected to indirect caliometry displayed increased oxygen consumption (VO_2_), carbon dioxide production (VCO_2_) and respiratory exchange ratio (RER), and decreased energy expenditure and physical activity, but no differences in food or water intake (**Figure 1F** and **Supplementary Figure 1B** and **C**). These results suggest that carbohydrate is the preferred energy source in PDH^SkM-/-^ mice due to its low fat content, but that is inadequate to support basal metabolic rates.

**Figure 1.**
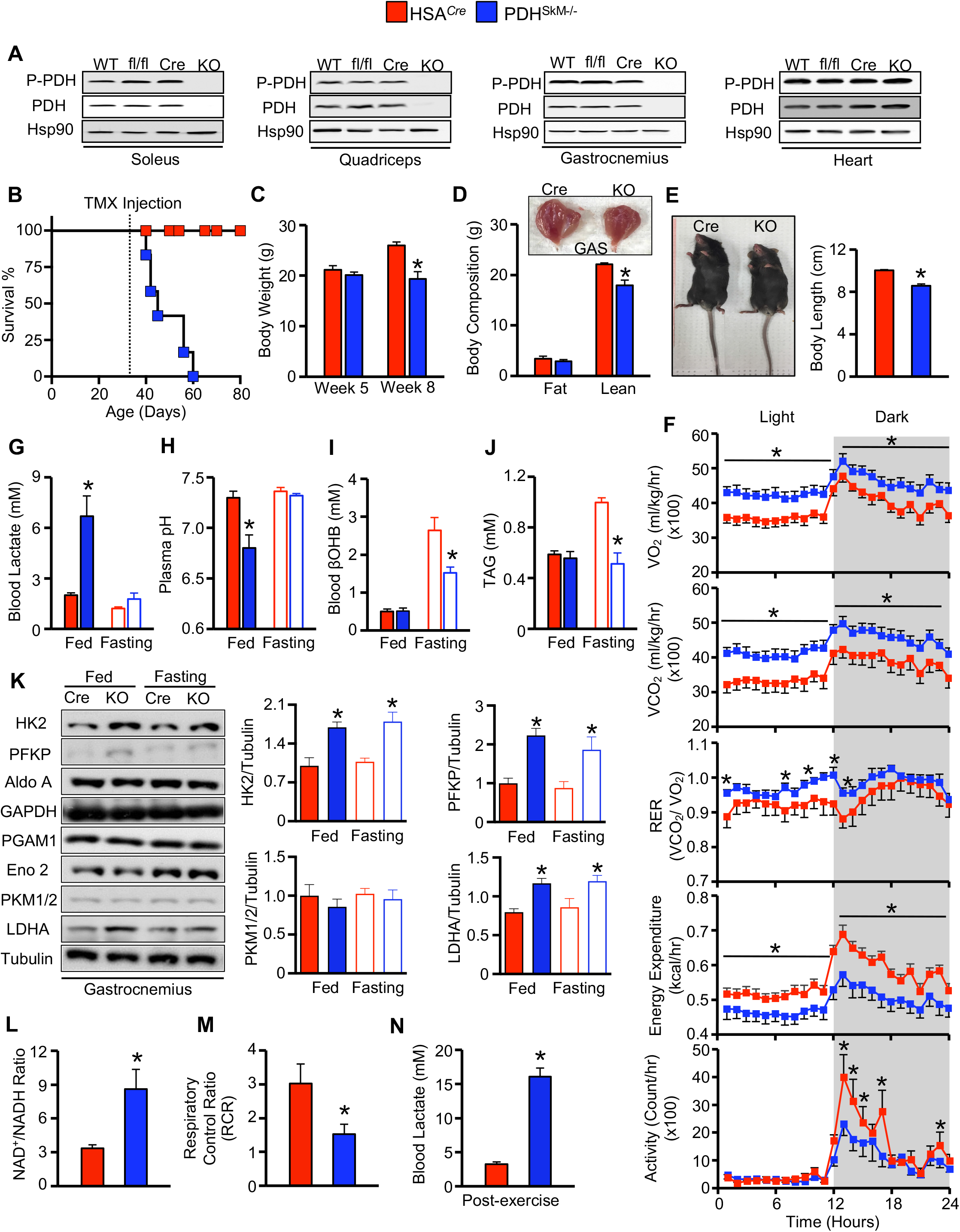
Skeletal muscle PDH deficiency induces lactic acidosis and premature mortality. (**A**) PDH protein expression in soleus, quadriceps, gastrocnemius, and heart in PDH^SkM-/-^ mice and their control littermates (fl/fl = floxed PDH littermate, HSA^*Cre*^ = human α-skeletal actin Cre expressing littermate, WT = wild-type) following tamoxifen injection (*n*=5-6). (**B-E**) Survival curve (**B**), body weight (**C**), body composition (**D**), and body length (**E**) were assessed in PDH^SkM-/-^ and HSA^*Cre*^ littermates post-tamoxifen injection (*n*=7-8). (**F**) Whole-body O_2_ consumption, CO_2_ production, respiratory exchange ratios (RER), energy expenditure, and ambulatory activity were measured for 24-hr after one day in the metabolic cages (*n*=5-6). (**G-K**) Levels of circulating lactate (**G**), pH (**H**), β-hydroxybutyrate (βOHB) (**I**), triacylglycerol (TAG) (**J**), and expression of muscle glycolytic enzymes (**K**) were assessed in *ad libitum* and fasting state (*n*=7-8). (**L-M**) NAD^+^/NADH ratio (**L**) and respiratory control ratio (RCR) (**M**) were measured in *ad libitum* state (*n*=6-7). (**N**) Blood lactate levels were measured post-exercise in PDH^SkM-/-^ and HSA^*Cre*^ littermates (*n*=6-7). Values are presented as means ± SEM. Statistical significance was determined using an unpaired two-tailed Student’s *t*-test or repeated measures ANOVA followed by a Bonferroni *post-hoc* analysis. \**P* < 0.05 vs. HSA^*Cre*^ mice.

Further blood biochemistry analysis revealed that PDH^SkM-/-^ mice suffered from robust lactic acidosis as their blood lactate concentrations were significantly elevated, coinciding with reduced blood pH levels under *ad libitum* feeding conditions (**Figure 1G** and **H**). Blood glucose concentrations were similar between the two groups (**Supplementary Figure 1D**), while concentrations of blood lipids, including ketone bodies and triacylglycerols (TAGs) were diminished in the PDH^SkM-/-^ mice during fasting (**Figure 1I** and **J**). To determine the causes that led to this deleterious lactic acidosis, we analyzed the glycolytic pathway in gastrocnemius muscles from HSA^*Cre*^ and PDH^SkM-/-^ mice. Muscle lacking PDH showed higher glycolytic protein expression, hexokinase 2 (HK2), platelet-type phosphofructokinase (PFKP), and lactate dehydrogenase A (LDHA), under both *ad libitum* and fasted conditions (**Figure 1K**). This observation was accompanied by higher glycolytic gene expression (e.g., *Gpd2, Ldha* and *Phka1*) and NAD^+^-to-NADH ratio (**Supplementary Figure 1E** and **Figure 1L**), and lower respiratory control ratio (RCR) levels in muscles from PDH^SkM-/-^ mice (**Figure 1M**). Most cytoplasmic pyruvate derived from glycolysis is either reduced to lactate by LDHA, oxidized to acetyl-CoA by PDH, or both. Accordingly, our results suggest that in the absence of glucose oxidation, a shift from oxidative to glycolytic metabolism leads to the accumulation of lactate. Next, we sought to determine whether lactic acidosis is the primary cause of this premature lethality in PDH^SkM-/-^ mice. To achieve this, approximately 1 week post-muscle PDH deletion, we subjected mice to a run-to-exhaustion protocol on a motorized treadmill. We observed that PDH^SkM-/-^ mice had extremely poor exercise tolerance, ran less than half the the distance and time as their HSA^*Cre*^ littermates, which led to an immediate decline in health status requiring the mice to be euthanized following the experiment (data not shown). Blood lactate concentrations post-exercise were elevated by approximately 400% in PDH^SkM-/-^ mice (**Figure 1N**). We concluded from these studies that the lethality of PDH^SkM-/-^ mice was likely induced by severe lactic acidosis.

### Loss of Muscle PDH Promotes Pyruvate-Alanine Cycling and Anaplerotic Glutaminolysis

Pyruvate is interchangeable with alanine by alanine transaminase enzyme, formally known as glutamate-pyruvate transaminase (GPT), that is expressed in both the cytosol and mitochondria. To examine the role of pyruvate-alanine cycling in the absence of muscle PDH, we performed targeted metabolomics analysis on gastrocnemius muscles collected from both genotypes during the *ad libitum* state. We observed that intramuscular alanine was increased by approximately 80% in PDH^SkM-/-^ mice compared to their HSA^*Cre*^ littermates (**Figure 2A**). qPCR analysis revealed that only mitochondrial *Gpt2* gene expression, but not cytoplasmic *Gpt1* gene expression, was elevated in the muscles of PDH^SkM-/-^ mice (**Figure 2B**), suggesting increased pyruvate transamination to alanine. Mitochondrial pyruvate can bypass PDH and be converted directly to oxaloacetate or malate by pyruvate carboxylase (PC) or malic enzyme (ME), respectively. mRNA expression of *Pc1* did not show differences between the two groups (**Supplementary Figure 1F**). In contrast, mRNA expression of *Me2* and *Me3* were significantly higher in the muscle of PDH^SkM-/-^ mice compared with HSA^*Cre*^ controls (**Figure 2C**). These data suggest that the absence of muscle PDH induces pyruvate-alanine cycling and triggers pyruvate-malate interconversions.

**Figure 2.**
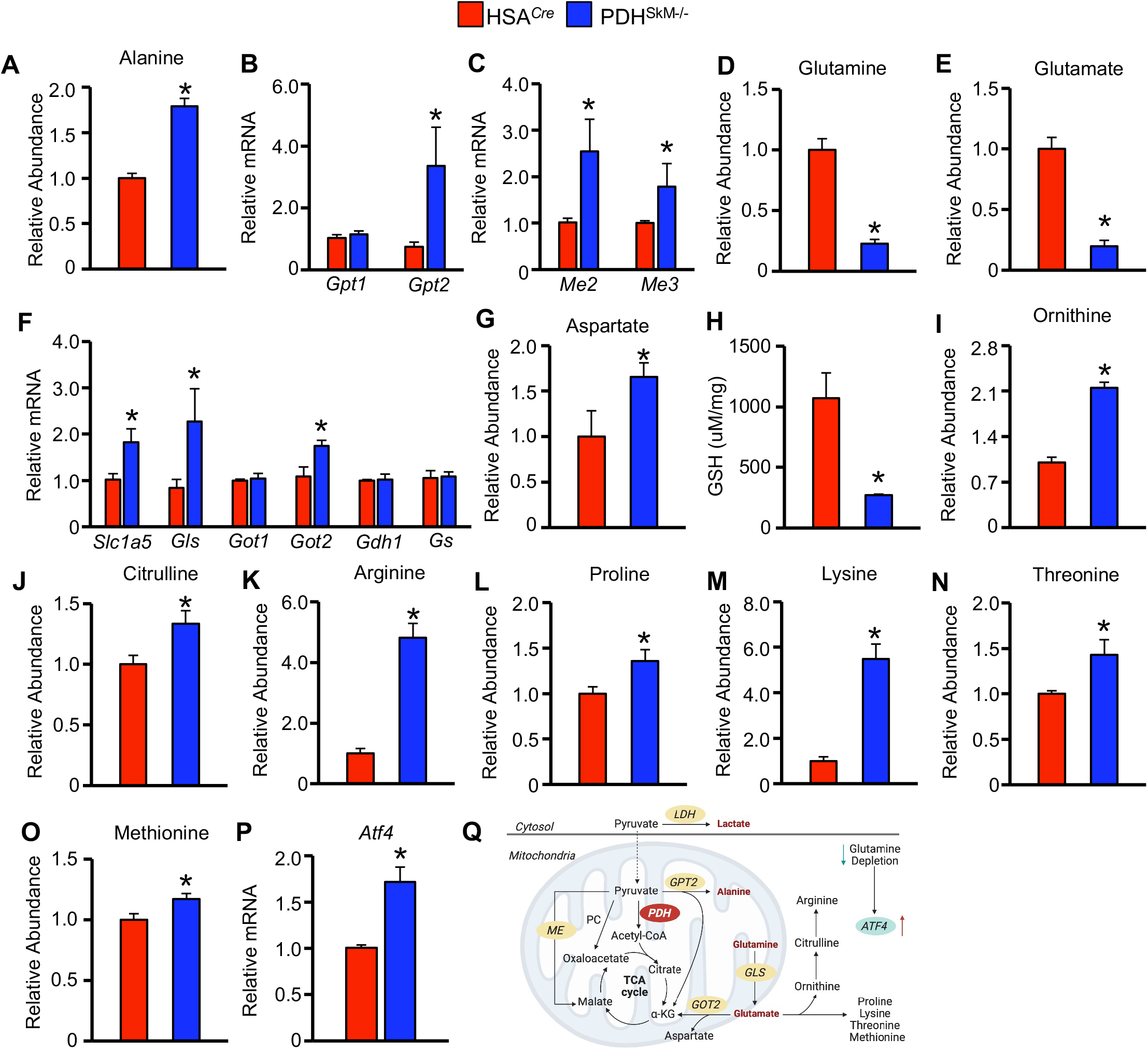
Skeletal muscle PDH deficiency promotes pyruvate-alanine cycling and glutaminolysis. (**A-P**) Levels of alanine (**A**), mRNA expression of glutamate-pyruvate transaminases (*Gpt*) 1 and 2, (**B**) and malic enzymes (*Me*) (**C**), glutamine (**D**), glutamate (**E**), mRNA expression of glutaminolysis genes (**F**), aspartate (**G**), glutathione (GSH) (**H**), ornithine (**I**), citrulline (**J**), arginine (**K**), proline (**L**), lysine (**M**), threonine (**N**), methionine (**O**), and mRNA expression of activating transcription factor (*Atf*) 4 (**P**) were measured in gastrocnemius muscles collected from PDH^SkM-/-^ and HSA^*Cre*^ littermates during the *ad libitum* state (*n*= 6-7). (**Q**) A schematic depicting the increased utilization of glutamine and pyruvate-alanine cycle activity induced by the loss of muscle PDH. Values are presented as means ± SEM. Statistical significance was determined using an unpaired two-tailed Student’s *t*-test. \**P* < 0.05 vs. HSA^*Cre*^ mice.

Another alteration from the metabolomics analysis that caught our attention was a marked decline in glutamine and glutamate levels in muscles from PDH^SkM-/-^ mice compared to their HSA^*Cre*^ littermates (**Figure 2D** and **E**). To further investigate the role of glutamine metabolism in muscle PDH deficiency, we measured the expression of various genes that regulates glutamine metabolism. Most glutaminolysis-related genes were highly expressed in muscle PDH^SkM-/-^ mice. Notably, expression of glutamine transporter (*Slc1a5*), glutaminase (*Gls*), and mitochondrial glutamic oxaloacetic transaminase 2 (*Got2*) were markedly higher in muscles from PDH^SkM-/-^ mice compared to those from HSA^*Cre*^ mice (**Figure 2F**). Nevertheless, we observed no changes in mRNA levels of glutamate dehydrogenase (GLUD or *Gdh1*), cytoplasmic *Got1*, and glutamine synthase (*Gs*) between the two groups (**Figure 2F**). In the glutaminolysis pathway, mitochondrial glutamate is converted to α⍰ketoglutarate (α-KG) by GLUD or aminotransferase (GPT2 and GOT2). GPT2 produces α-KG and alanine and couples the pyruvate-alanine cycle with glutaminolysis, while GOT2 produces α-KG and aspartate^10^. To gain further insight into glutaminolysis, muscle aspartate levels were measured in both groups. As expected, intramuscular aspartate was significantly increased in PDH^SkM-/-^ mice (**Figure 2G**). Another fate of mitochondrial glutamine-derived glutamate is the production of glutathione (GSH) and the synthesis of non-essential amino acids (NEAAs)^11^. Remarkably, muscle GSH levels were strikingly decreased in PDH^SkM-/-^ mice compared to their HSA^*Cre*^ littermates (**Figure 2H**). Conversely, intramuscular NEAA levels, including ornithine, citrulline, arginine, proline, lysine, threonine, and methionine, were significantly elevated in PDH^SkM-/-^ mice (**Figure 2I-O**). Of interest, the elevation in muscle NEAA levels and glutamine depletion of PDH^SkM-/-^ mice was coupled with marked increases in mRNA expression of activating transcription factor 4 (*Atf4*) (**Figure 2P**), a master transcriptional regulator stimulated under stress conditions and glutamine deprivation^12^. Together, these results suggest that impairing muscle PDH redirects glutamine away from GSH production toward NEAA synthesis and replenishing tricarboxylic acid (TCA) cycle intermediates.

Since muscles oxidize essential amino acids, including branched-chain amino acids (BCAAs), as an alternative substrate to glucose, we quantified BCAAs levels in gastrocnemius muscles collected from PDH^SkM-/-^ and HSA^*Cre*^ mice during the *ad libitum* state, and saw no changes between the two groups (**Supplementary Figure 1G**). We also examined muscle dependency on lipid metabolism and saw no significant differences in gene expression of key lipolytic markers (*Lpl, Pnpla2, Lipe*), fatty acid transporters (*Cd36, Cpt1*α, *Cpt1*β), and fatty acid binding and oxidation markers (*Fabp3, Acads, Acadm, Acadl, Hadh*) in muscles of PDH^SkM-/-^ mice versus HSA^*Cre*^ mice (**Supplementary Figure 1H-K**). Collectively, these results provided further evidence that loss of muscle PDH induces adaptive utilization of glutamine and increased pyruvate-alanine cycle activities (**Figure 2Q**).

### High-Fat Diet Supplementation Rescues the Metabolic Defects Induced by Muscle PDH Deficiency

Next, we reasoned that PDH^SkM-/-^ mice did not survive on a LFD due to the dearth of glucose oxidation and the lack of fatty acid availability. To investigate this possibility, we subjected HSA^*Cre*^ and PDH^SkM-/-^ mice to high-fat diet (HFD) supplementation for 10 weeks. Surprisingly, PDH^SkM-/-^ mice lived and gained weight at equal rates to HSA^*Cre*^ littermates (**Figure 3A**). No mortality was recorded during the period of the study. Both groups had similar lean mass and total body length; however, PDH^SkM-/-^ mice showed slightly but not significantly lower fat mass (**Figure 3B** and **C**). Blood glucose concentrations were comparable between the groups under *ad libitum* and fasting conditions (**Figure 3D**). Likewise, no significant differences were detected in circulating ketone bodies, TAGs, non-esterified fatty acids and cholesterol between the groups (**Supplementary Figure 2A-D**). Interestingly, *ad libitum* blood lactate concentrations were higher in PDH^SkM-/-^ mice with no changes in pH levels (**Figure 3E** and **Supplementary Figure 2E**), indicating that when fed a HFD, PDH^SkM-/-^ mice exhibited hyperlactatemia but not lactic acidosis. Analysis of the glycolytic pathway in gastrocnemius muscles by qPCR and western blotting revealed that increased lactate production in PDH^SkM-/-^ mice was unlikely a result of enhanced glycolysis (**Supplementary Figure 2F** and **G**). We further confirmed these results by measuring NAD^+^, NADH and NAD^+^-to-NADH ratio in quadriceps muscles, and we saw no differences among the two groups (**Supplementary Figure 2H-J**).

**Figure 3.**
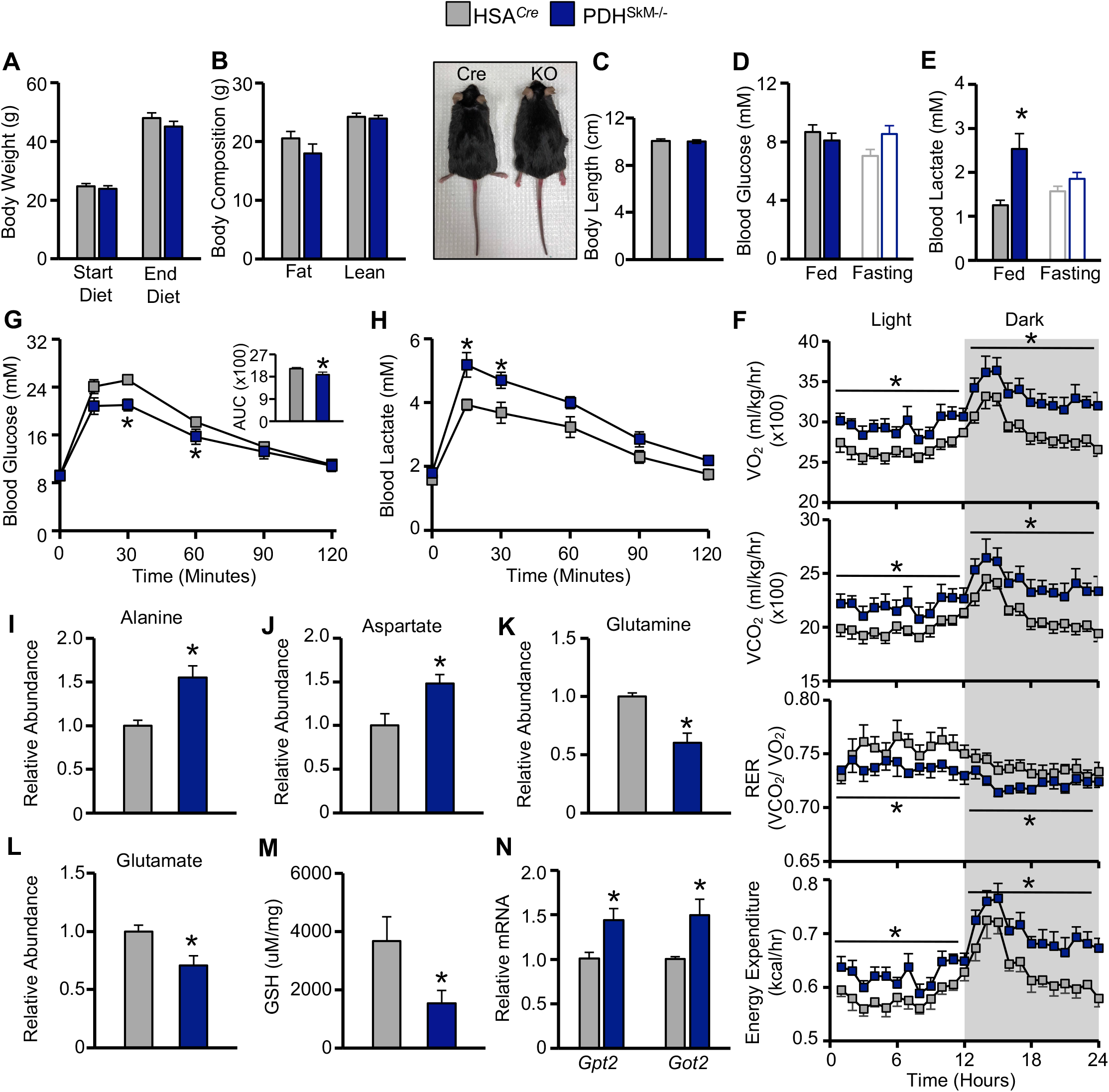
HFD supplementation abolishes death and overt phenotype induced by muscle PDH deficiency. (**A-E**) Body weight (**A**), body composition (**B**), body length (**C**), blood glucose (**D**), and blood lactate (**E**) were measured in obese PDH^SkM-/-^ and HSA^*Cre*^ mice (*n*=7-8). (**F**) Whole-body O_2_ consumption, CO_2_ production, respiratory exchange ratios (RER), and energy expenditure were measured for 24-hr after one day in the metabolic cages (*n*=5-6). (**G and H**) Glucose tolerance (**G**) and lactate (**H**) levels were detected at different timepoints during the intraperitoneal glucose tolerance test (IP-GTT) in obese PDH^SkM-/-^ and HSA^*Cre*^ mice (*n*=9-10). (**I-N**) Levels of alanine (**I**), aspartate (**J**), glutamine (**K**), glutamate (**L**), glutathione (GSH) (**M**), and mRNA expression of glutamate-pyruvate transaminases (*Gpt*) 2 and glutamic oxaloacetic transaminase (*Got*) 2 (**N**) were evaluated in gastrocnemius muscles collected from obese PDH^SkM-/-^ and HSA^*Cre*^ mice during the *ad libitum* state (*n*=5-6). Values are presented as means ± SEM. Statistical significance was determined using an unpaired two-tailed Student’s *t*-test or repeated measures ANOVA followed by a Bonferroni *post-hoc* analysis. \**P* < 0.05 vs. HSA^*Cre*^ mice.

To examine whether loss of muscle PDH induces alterations in whole-body metabolism during HFD supplementation, we performed metabolic chamber studies in HSA^*Cre*^ and PDH^SkM-/-^ mice fed an HFD. In contrast to when fed a LFD, PDH^SkM-/-^ mice manifested an increased VO_2_, VCO_2_, energy expenditure, and decreased RER (**Figure 3F**), suggesting a substrate shift in favor of fatty acid oxidation. Of note, the difference in energy expenditure was conserved when accounting for any differences in body mass by ANCOVA (data not shown). As there were no changes in physical activity, food and water intake between the groups (**Supplementary Figure 2K-M**), we asked whether enhanced energy expenditure of PDH^SkM-/-^ obese mice could be attributed to increased thermogenesis. To this end, we saw a dramatic increase in gene expression of *Ucp1* and other thermogenesis-related genes (*Pgc-1*α, *Cidea, Cox8b*) in adipose tissue of PDH^SkM-/-^ mice compared to HSA^*Cre*^ mice (**Supplementary Figure 2N**). Furthermore, PDH^SkM-/-^ mice demonstrated improved intraperitoneal glucose tolerance (**Figure 3G**) without changes in insulin tolerance (**Supplementary Figure 2O**), implying that differences in glucose tolerance were not a result of improved peripheral insulin sensitivity. The improvement in glucose handling prompted us to explore further the mechanism underlying the phenotype of PDH^SkM-/-^ mice. HSA^*Cre*^ and PDH^SkM-/-^ mice fed an HFD underwent another glucose tolerance test, but we measured circulating lactate levels as a marker of lactate excursion. PDH^SkM-/-^ mice clearly showed higher lactate levels throughout the test than their control mice (**Figure 3H**), suggesting that the majority of administered glucose was converted into lactate. These results demonstrate that HFD supplementation rescued the phenotype in PDH^SkM-/-^ mice and increased fatty acid utilization to generate energy.

### Loss of Muscle PDH Promotes Pyruvate-Alanine Cycling and Glutaminolysis Utilization Even During HFD Supplementation

To further examine pyruvate-alanine cycling and glutamine metabolism in PDH^SkM-/-^ mice during HFD supplementation, we extended our targeted metabolomic analysis on gastrocnemius muscles harvested from HSA^*Cre*^ and PDH^SkM^ obese mice under *ad libitum* conditions. Similar to LFD provision, analysis of numerous metabolites showed apparent alterations in the abundance of several amino acids related to pyruvate transamination to alanine and glutaminolysis. Alanine and aspartate levels were significantly increased in muscle of PDH^SkM^ obese mice compared with their controls (**Figure 3I** and **J**). Interestingly, we observed no changes in muscle citrulline, ornithine and arginine levels between genotypes (**Supplementary Figure 2P**). However, this was not surprising because the magnitude of increased muscle alanine in PDH^SkM^ mice under HFD supplementation was less than what we observed during LFD provision. Additionally, both glutamine and glutamate levels were significantly reduced in PDH^SkM^ obese mice muscle compared with HSA^*Cre*^ control mice (**Figure 3K** and **L**), suggesting increased glutamine flux into the TCA cycle. Consistent with these observations, we detected significant decreases in intramuscular GSH and increases in muscle mRNA levels of *Gpt2* and *Got2* in PDH^SkM-/-^ obese mice (**Figure 2M** and **N**). Lastly, measurement of other essential amino acids (i.e., BCAAs) in gastrocnemius muscles revealed no significant differences between groups (**Supplementary Figure 2Q**). Taken together, these results suggest that loss of muscle PDH promotes pyruvate-alanine cycling and glutaminolysis even during HFD supplementation.

### Loss of Muscle PDH Compromises Exercise Performance Despite Increased Lipid Oxidation Under HFD Supplementation

To further explore the functional importance of aerobic glucose oxidation during exercise, we subjected HSA^*Cre*^ and PDH^SkM-/-^ obese mice to a low-intensity run to exhaustion treadmill running protocol. Relative to the performance of HSA^*Cre*^ controls, PDH^SkM-/-^ obese mice ran approximately 70 to 80 % less the time and distance (**Figure 4A** and **B**). Furthermore, this deficit in exercise performance in PDH^SkM-/-^ obese mice coincided with strikingly higher post-exercise blood lactate and lower pH levels (**Figure 4C** and **D**), but no changes in post-exercise blood glucose concentrations (**Figure 4E**). Notably however, no mortality was recorded in PDH^SkM-/-^ obese mice post-exercise. Because fatty acids are the primary energy source in skeletal muscle during rest and mild-intensity exercise, we examined muscle acylcarnitines as a marker for lipid oxidation using gastrocnemius muscles collected from both genotypes during sedentary and exercise conditions. Although global muscle acylcarnitine quantitative profile was similar in both groups under sedentary and exercise conditions (**Figure 4F**), short-, but not medium- or long-chain acylcarnitine species, were significantly decreased in muscles from PDH^SkM-/-^ obese mice compared against those from HSA^*Cre*^ controls during both sedentary and exercise conditions (**Figure 4G-I**), suggestive of increased muscle fatty acids utilization. Likewise, we observed significant increases in muscle carnitine acetyltransferase (CrAT) activity, a muscle-enriched enzyme that buffers the mitochondrial acetyl-CoA pool by converting short chain acyl-CoAs to their membrane permeant acylcarnitine analogues, in PDH^SkM-/-^ obese mice compared to their HSA^*Cre*^ controls during sedentary and exercise conditions (**Figure 4J**).

**Figure 4.**
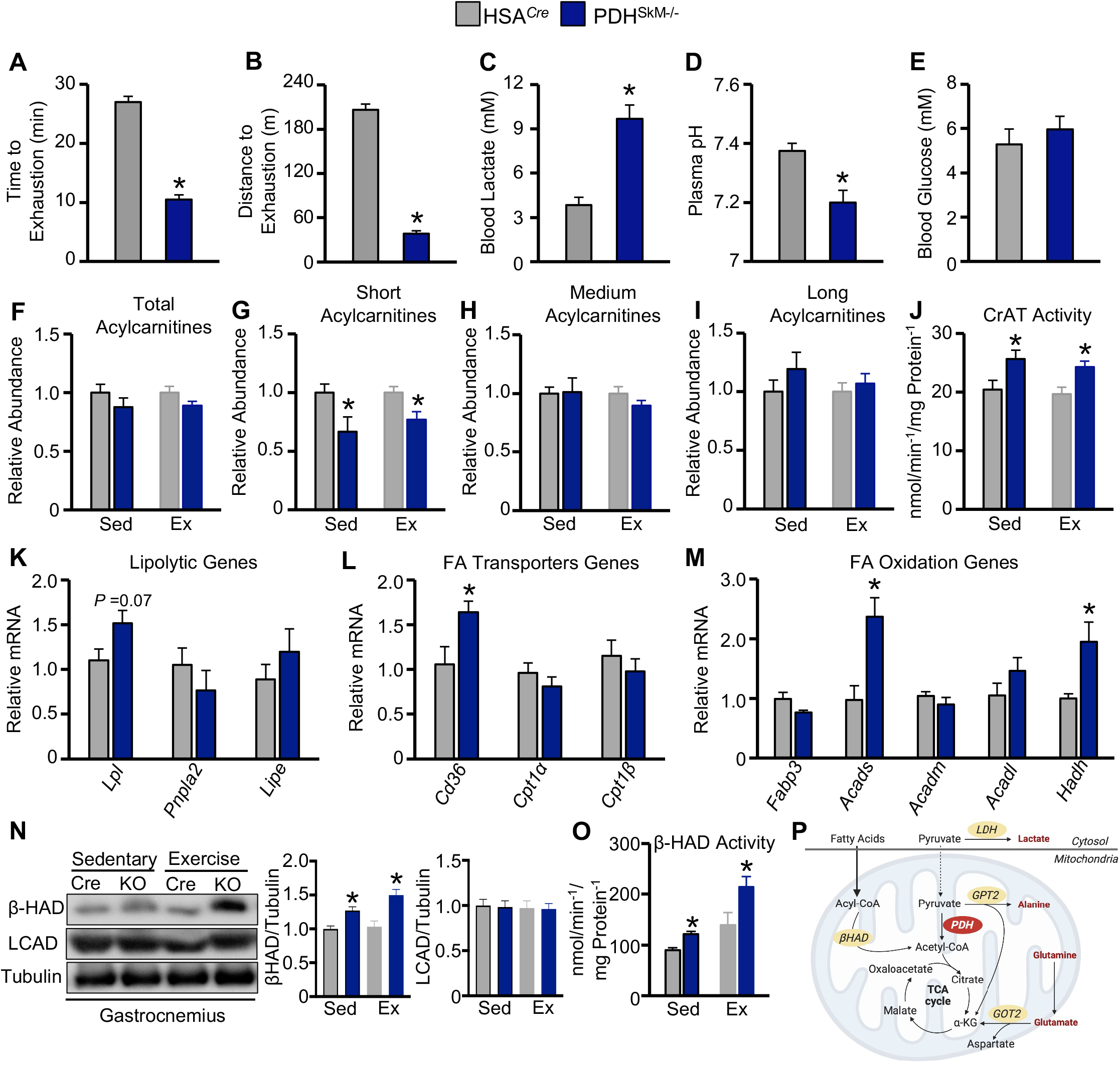
Skeletal muscle PDH deficiency compromised exercise performance despite increased fat oxidation under HFD supplementation. (**A-E**) Time to exhaustion (**A**), distance to exhaustion (**B**), blood lactate (**C**), plasma pH (**D**), and blood glucose (**E**) were evaluated during the exercise tolerance test in obese PDH^SkM-/-^ and HSA^*Cre*^ mice (*n*=6-7). (**F-O**) Total acylcarnitines (**F**), short acylcarnitines (**G**), medium acylcarnitines (**H**), long acylcarnitines (**I**), carnitine acetyltransferase (CrAT) activity (**J**), mRNA expression of lipolytic genes (**K**), fatty acids (FA) transporter genes (**L**) and oxidation genes (**M**), protein expression of fatty acid oxidation enzymes (long-chain acyl-CoA dehydrogenase (LCAD) and β-hydroxyacyl CoA dehydrogenase (β-HAD)) (**N**), and β-HAD activity (**O**) were evaluated in gastrocnemius muscles collected from obese PDH^SkM-/-^ and HSA^*Cre*^ mice during sedentary (Sed) or exercise (Ex) conditions (*n*=5-6). (**P**) A schematic of fatty acid and glucose metabolism in PDH^SkM-/-^ obese mice under sedentary and exercise conditions. Values are presented as means ± SEM. Statistical significance was determined using an unpaired two-tailed Student’s *t*-test. \**P* < 0.05 vs. HSA^*Cre*^ mice.

To gain more insight into muscle reliance on lipid metabolism during HFD supplementation, we analyzed the expression of genes involved in fatty acid catabolism in gastrocnemius muscles extracted from both groups under sedentary conditions. Interestingly, levels of mRNA encoding enzymes involved in lipolysis (*Lpl*), fatty acid uptake (*Cd36*) and short-chain fatty acid oxidation (*Acads, Hadh*) were augmented in muscle of PDH^SkM-/-^ obese mice when compared to those from HSA^*Cre*^ controls, whereas levels of mRNA encoding other enzymes involved in controlling medium- and long-chain fatty acid oxidation (*Acadm, Acadl*) were similar among the groups (**Figure 4K-M**). Western blot analysis of muscle lysates showed that only protein levels of β-hydroxyacyl CoA dehydrogenase (β-HAD), but not long-chain acyl CoA dehydrogenase (LCAD), were increased in PDH^SkM-/-^ obese mice under both sedentary and exercise conditions (**Figure 4N**). The augmented protein levels of β-HAD were mirrored by increased activity of β-HAD in muscle from PDH^SkM-/-^ obese mice under both conditions (**Figure 4O**). Taken together, these results suggest that fatty acid oxidation is enhanced in muscle PDH^SkM-/-^ obese mice under sedentary and exercise conditions (**Figure 4P**). Nevertheless, in the absence of muscle glucose oxidation, fatty acid oxidation alone is insufficient to meet high energy demand during exercise.

## DISCUSSION

PDH is a part of the mitochondrial PDH complex that catalyzes the oxidative decarboxylation of pyruvate to form acetyl-CoA, and links glucose metabolism to the mitochondrial TCA cycle. To explore the significance of PDH in metabolic flexibility, PDH was deliberately removed from skeletal muscle, which is a valuable model for studying metabolic flexibility due to its ability to adapt fuel utilization based on the available fuel sources^13^. Results described herein provide compelling evidence regarding the importance of muscle PDH in determining the fate of pyruvate produced from glycolysis, metabolic reprogramming, and survival. Our findings demonstrate that muscle PDH deletion substantially augments the glycolytic pathway and rewires the generated cytosolic pyruvate away from glycose oxidation to towards lactate production by LDH, a phenomenon that induces severe lactic acidosis and causes premature death at rest and sudden death post-low-intensity exercise in mice lacking muscle PDH under LFD provision. These results are consistent with a previous study that showed that mice with a muscle- and heart-specific PDH deficiency die within 7 days of weaning when fed a chow diet^9^, but inconsistent with a recent study that demonstrated that muscle PDH ablation in adult mice fed a chow diet exhibited no overt phenotype and survived premature mortality^14^. The discrepancies between the latter study and our results could be attributable to the differences in micronutrients and macronutrients and the magnitude of muscle PDH deletion between the two studies. Furthermore, It is possible that the developmental stage at which PDH ablation occurs could affect the severity of the phenotype observed in the two studies.

The mitochondrial pyruvate carrier (MPC) is a protein complex that plays a crucial role in importing pyruvate into the mitochondria^15,16^. Once inside the mitochondria, pyruvate is converted directly to acetyl-CoA by PDH and fuels the TCA cycle or is transformed into various TCA cycle intermediates through anaplerotic reactions^17^. The present study reveals that the depletion of muscle PDH prompts an anaplerotic response by augmenting pyruvate-alanine transamination and pyruvate carboxylation to malate mediated by GPT2 and ME, respectively. Furthermore, our observations demonstrate that muscle PDH deficiency increases muscle reliance on glutamine catabolism and diverts glutamine from GSH biosynthesis towards α-KG to replenish the TCA cycle. The substantial rise in glutaminolysis resulting from muscle PDH deficiency amplifies the biosynthesis of NEAAs from glutamate derived from glutamine. Our results are consistent with earlier investigations indicating that impeding the entry of pyruvate into the mitochondria through the deletion of MPC1 or MPC2 in the skeletal muscle or liver elevates pyruvate-alanine cycling and augments glutaminolysis as an adaptive mechanism to replenish the TCA cycle^18-21^. Despite the observed elevation of anaplerotic reactions, muscle PDH deficiency curtails the muscle’s metabolic flexibility to switch toward lipid catabolism during LFD provision. Consequently, this leads to a rapid decline in body weight and muscle mass and causes premature mortality following PDH inactivation when fatty acid availability is low.

In contrast to what was seen with LFD provision, mice lacking muscle PDH that were fed an HFD survived without displaying any noticeable phenotype. Body composition and length remained unchanged between the two genotypes throughout the study. Notably, even under HFD supplementation, muscle PDH deficiency induces hyperlactatemia without any concomitant changes in blood pH levels and glucose concentrations. Surprisingly, obese mice lacking muscle PDH exhibited improvement in whole-body glucose handling, without any discernable changes in insulin sensitivity. This finding may appear counterintuitive because it is well-known that diminished muscle glucose oxidation during obesity is associated with impaired glucose tolerance and deteriorated insulin sensitivity^22,23^. In support of this, previous study have shown that mice with whole-body deletion of PDK4, a mitochondrial enzyme that inhibits the activity of PDH, are protected against insulin resistance when fed an HFD^24^. Nevertheless, further analysis revealed that the reason for the observed protection in our knockout mice during HFD supplementation is a result of increased muscle lactate excretion, possibly to offset and balance glucose uptake with disposal. As such, our findings provide evidence that impairments in muscle PDH activity increase glucose disposal by directly converting glucose-derived pyruvate into lactate, thereby improving glucose tolerance. However, further investigations are necessary to comprehensively delineate the ultimate fate of pyruvate produced by glycolysis.

Another important finding that emerges from this study is that the disruption of muscle PDH augments the reliance on fatty acid oxidation under HFD supplementation. At the whole-body level, we observed that muscle PDH deficiency leads to an increase in oxygen consumption and a decrease in RER during the light and dark cycle, suggesting an increased reliance on fatty acid oxidation as an adaptive measure to compensate for decreased glucose oxidation. At the cellular levels, our results demonstrate that muscle PDH deficiency stimulates fatty acid uptake and oxidation. These findings are further supported by a reduction in muscle short acylcarnitine levels, coupled with increased activity of CrAT, which selectively transports short-chain fatty acyl-CoAs into the mitochondrial matrix for β-oxidation to ensue^25,26^. Interestingly, despite evidence of increased fatty acid oxidation and CrAT activity under HFD supplementation, obese mice with disrupted muscle PDH showed impairment in exercise performance and flexibility to fully switch to glucose oxidation during higher energy demand, leading to early fatigue. We therefore surmise that deficits in muscle PDH activity induce metabolic inflexibility that manifests when there is a rapid switch between glucose and fatty acid oxidation during the transition from rest to exercise. These findings lend merit to the glucose-fatty acid cycle originally described by Randle and colleagues^2^ and the concept of metabolic inflexibility proposed by Kelley and Mandarino^3^.

In conclusion, the present study demonstrates the crucial role of PDH in metabolic flexibility and the consequences of its removal from the skeletal muscle. Our results reveal that elimination of PDH in the skeletal muscle elevates dependence on glutaminolysis, augments pyruvate-alanine transamination and pyruvate carboxylation, and diverts glutamine from GSH biosynthesis to replenish the TCA cycle. In addition, PDH deletion also curtails muscle metabolic flexibility and impairs its ability to switch to lipid catabolism during a LFD provision, leading to premature mortality. Ironically, mice lacking muscle PDH that were fed an HFD survived longer, mainly due to increase in reliance on fatty acid oxidation; however, their exercise performance was compromised.

## SUPPLEMENTAL INFORMATION

Supplemental information is available online.

## ACKNOWLEDGEMENTS & FUNDING

All the schematics in this study were created with BioRender.com. This work was supported by the Natural Sciences and Engineering Research Council (to R.A.B.), the Canada Foundation for Innovation (to R.A.B.), and Canadian Institutes of Health Research (to J.R.U.). J.R.U. is a Tier 2 Canada Research Chair (Pharmacotherapy of Energy Metabolism in Obesity), and R.A.B. is a Research Scholar of the Fonds de Recherche du Québec - Santé (FRQS) and a New Investigator of the Kidney Research Scientist Core Education and National Training (KRESCENT).

## AUTHOR CONTRIBUTIONS

Conceptualization, K.G., A.M.A., and R.A.B.; Investigation, K.G., A.M.A., X.L., A.A.G., Q.G.K., C.S., G.M.U., A.M.D., K.L.J., T.R.A., and F.E.; Formal Analysis and Visualization, K.G., and A.M.A.; Writing – Original Draft, K.G., A.M.A., and R.A.B.; Writing – Review & Editing, K.G., A.M.A., J.M.S., G.D.L., J.R.U., and R.A.B.; Funding Acquisition and Project Administration, R.A.B. and J.R.U.; Resources, J.M.S., G.D.L., J.R.U., and R.A.B.; Supervision, R.A.B.; R.A.B. takes full responsibility for the data within this paper.

## FIGURE LEGENDS

**Supplementary Figure S1. Skeletal muscle PDH deficiency induces lactic acidosis, pyruvate-alanine cycling, and glutaminolysis. Related to Figures 1 and 2**.

(**A**) mRNA expression of genes that regulate the PDH complex in gastrocnemius muscles collected from PDH^SkM-/-^ and HSA^*Cre*^ littermates during the *ad libitum* state (*n*=5-6). (**B-D**) Food intake (**B**), water intake (**C**), and blood glucose (**D**) were measured during the *ad libitum* state (*n*=5-6). (**E-J**) mRNA expression of glucose metabolism (**E**) and pyruvate carboxylase (*Pc*) 1 (**F**) genes, levels of branched-chain amino acids (BCAAs) (**G**), and mRNA expression of lipolytic (**H**), fatty acid transporters (**I**) and oxidation (**J**) genes were evaluated in gastrocnemius muscles of PDH^SkM-/-^ and HSA^*Cre*^ mice (*n*=5-6). Values are presented as means ± SEM. Statistical significance was determined using an unpaired two-tailed Student’s *t*-test. \**P* < 0.05 vs. HSA^*Cre*^ mice.

**Supplementary Figure S2. HFD supplementation abolishes overt phenotype induced by muscle PDH deficiency. Related to Figure 3 and 4**.

(**A-E**) Levels of circulating β-hydroxybutyrate (βOHB) (**A**), triacylglycerol (TAG) (**B**), non-esterified fatty acids (NEFA) (**C**), cholesterol (**D**), and pH (**E**) were measured in obese PDH^SkM-/-^ and HSA^*Cre*^ mice (*n*= 6-7). (**F-J**) mRNA (**F**) and protein (**G**) expression of glucose metabolism regulators, NAD+ levels (**H**), NADH level (**I**), and NAD^+^/NADH ratio (**J**) were evaluated in gastrocnemius muscles of obese PDH^SkM-/-^ and HSA^*Cre*^ mice (*n*=5-6). (**K-M**) Ambulatory activity (**K**), food intake (**L**), and water intake (**M**) were measured for 24-hr after one day in the metabolic cages (*n*=5-6). (**N-Q**) mRNA expression of thermogenesis-related genes in adipose tissues (**N**), insulin tolerance (**O**), and levels of citrulline/ornithine/arginine (**P**) and branched-chain amino acids (BCAAs) (**Q**) were measured in gastrocnemius muscles of obese PDH^SkM-/-^ and HSA^*Cre*^ mice (*n*= 6-7). Values are presented as means ± SEM. Statistical significance was determined using an unpaired two-tailed Student’s *t*-test or repeated measures ANOVA followed by a Bonferroni *post-hoc* analysis. \**P* < 0.05 vs. HSA^*Cre*^ mice.

## STAR⋆ METHODS

### LEAD CONTACT AND MATERIALS AVAILABILITY

Further information and requests for resources and reagents should be directed to the Lead Contact for this manuscript: Rami Al Batran.

### EXPERIMENTAL MODELS

#### Animal care and experimentation

##### Generation of skeletal muscle-specific PDH^SkM-/-^ mice

Skeletal muscle-specific PDH^SkM-/-^ mice were generated by crossing PDH^Flox^ mice with HSA^*Cre*^ mice (The Jackson Laboratory), which express tamoxifen-inducible Cre recombinase under the control of the skeletal muscle-specific HSA-Cre promoter. To induce skeletal muscle-specific PDH deletion, 5-week-old male PDH^SkM-/-^ mice and HSA^*Cre*^ littermates were intraperitoneally (IP) injected with tamoxifen (50 mg/kg) dissolved in corn oil for 5 consecutive days, as we have previously described^27^. Tamoxifen administration activates Cre recombinase specifically in skeletal muscle cells, leading to excision of the loxP-flanked DNA regions of PDH. After the end of the experimental protocol, PDH^SkM-/-^ mice and HSA^*Cre*^ littermates were euthanized, tissues were extracted and immediately snap-frozen in liquid N_2_ using liquid N_2_-cooled Wollenberger tongs. This ensured that the samples were preserved in a manner that allowed for subsequent biochemical analyses.

##### Animal care

All animal procedures were performed in accordance with the guidelines of the Canadian Council on Animal Care and approved by the institute’s Health Sciences Animal Welfare Committee. Male HSA^*Cre*^ and PDH^SkM-/-^ mice aged 7 weeks were fed either a low-fat diet (10% kcal from lard, D12450J) or a high-fat diet (60% kcal from lard, Research Diets; D12492) for 10-weeks to induce obesity. At the end of the study, animals were euthanized via an IP injection of sodium pentobarbital (12 mg) while in either the in the ad libitum or fasted state. Tissues, such as gastrocnemius muscles and liver, were immediately collected and snap-frozen in liquid N_2_ using liquid N_2_-cooled Wollenberger tongs.

##### Assessment of glucose homeostasis

Glucose and insulin tolerance tests were performed in overnight-fasted mice. Mice were injected intraperitoneally with glucose (2 g/kg) or insulin (0.5 U/kg). Blood glucose levels were measured at 0, 15, 30, 60, 90, and 120 minutes after glucose or insulin administration using the Contour Next blood glucose monitoring system (Bayer®). Whole-blood samples were collected from the tail vein of each mouse for analysis.

##### Magnetic resonance imaging

Body composition analysis of mice was performed using quantitative nuclear magnetic resonance relaxometry. The EchoMRI-body composition analyzer was utilized for quantification of total lean and fat mass as previously described^28^.

##### *In Vivo* metabolic assessment

*In vivo* metabolic assessments were performed using the Oxymax laboratory animal monitoring system (Columbus Instruments), which utilizes indirect calorimetry. Prior to data collection, mice were acclimatized to the system for a 24-hour period, as previously described^28^. During the metabolic assessment, various parameters were quantified, including animal activity, food intake, water intake, whole-body oxygen consumption rates, and respiratory exchange ratios. These measurements provided valuable insights into the metabolic state of the animals and allowed for the characterization of changes in energy expenditure and substrate utilization under different conditions.

##### Exercise Capacity Testing

Exercise capacity was performed by running mice on a calibrated, motor-driven treadmill (Columbus Instruments) at a speed of 3 m/min for 1 min, followed by a speed increase of 4 m/min for 1 min, 5 m/min for 1 min, 6 m/min for 3-min, 8 m/min for 14-min, 9 m/min for 10-min, 10 m/min for 7-min, 12 m/min for 7-min, and 14 m/min until exhaustion as previously described^29^. The data were collected after the first 6 min of the acclimatization period. Exhaustion was determined as the mice spending >10 consecutive seconds on the shock grid or running off the shock grid and immediately falling back onto the shock grid 3 consecutive times.

### METHOD DETAILS

#### Blood chemistry analysis

Blood glucose levels were measured in tail whole-blood samples obtained during the random fed state or following a 16-hour fast using the Contour Next blood glucose monitoring system (Bayer®). βOHB levels were measured using the FreeStyle Precision Blood β-ketone meter (Abbott Laboratories), while lactate levels were measured using the Lactate Plus Meter (Nova Biomedical). Plasma samples were collected from mice and analyzed using an EasyRA® clinical chemistry analyzer (Medica, Bedford, USA) to determine the levels of alanine aminotransferase, aspartate aminotransferase, albumin, and cholesterol. This allowed for the quantification of key biomarkers related to liver function, lipid metabolism, and protein metabolism, providing valuable information about the physiological status of the animals.

#### Western blotting

Frozen tissue samples (20 mg) were homogenized using a buffer containing 50 mM Tris-HCl (pH 8 at 4°C), 1 mM EDTA, 10% glycerol (w/v), 0.02% Brij-35 (w/v), 1 mM dithiothreitol (DTT), and a cocktail of protease and phosphatase inhibitors (Sigma). Protein samples were then prepared and subjected to western blotting protocols, as previously described^30^.

#### Real-time PCR analysis

Total RNA was extracted from the tissue samples using a commercial kit according to the manufacturer’s instructions. First-strand cDNA synthesis was performed using a high-capacity cDNA reverse transcription kit (Applied Biosystems). Real-time PCR was carried out using SYBR green (Kapa Biosystems, Inc.) and the CFX connect real-time PCR machine (Bio-Rad Laboratories Inc.). The quantification of relative mRNA transcript levels was performed using the 2^-ΔΔCt^ method^31^, with the peptidylprolyl isomerase A (*Ppia*) gene serving as our internal control for normalization.

#### Determination of triacylglycerols content

The frozen gastrocnemius tissues (∼20 mg) were homogenized in a 2:1 chloroform:methanol solution, and the resulting supernatant was used for the determination of triacylglycerols (TAGs) content using an enzymatic assay kit (Wako Pure Chemical Industries), as previously described^32^. The same enzymatic assay kit was used to measure TAG levels in mouse plasma samples (4 μL), as previously described^32^.

#### Metabolomic profiling

To perform targeted quantitative metabolomics, frozen powdered gastrocnemius tissues (∼50 mg) were subjected to direct injection mass spectrometry (DI-MS) combined with reverse-phase liquid chromatography (LC)-MS/MS assay as previously described^33^. Isotope-labeled internal standards were used for metabolite quantification. Data analysis was performed and concentrations were calculated using Analyst software.

#### Assessment of glutathione levels

Gastrocnemius muscle samples were used to measure glutathione (GSH) levels via a fluorometric assay (Abcam, Ab138881) according to the manufacturer’s protocol. Briefly, frozen tissue samples were lysed in 1% NP-40 and subsequently diluted 10-fold in reaction buffer. Fluorescence was measured for 10-40 minutes after the addition of dye, as previously described^34^.

#### Carnitine acetyltransferase activity

Carnitine acetyltransferase (CrAT) activity was measured in gastrocnemius muscle samples according to previously described methods^35^. Briefly, the reaction mixture consisted of 0.1 M Tris-HCl (pH 8.0), 125 μM 5,5’-Dithiobis-(2-Nitrobenzoic Acid) (DTNB), 0.1 mM acetyl-CoA, 1.1 mM L-carnitine, and an aliquot of the enzyme source with a standardized protein concentration of mg/ml in triplicate wells of a 96-well plate. The reaction produces acetyl-carnitine and free coenzyme A (CoA), which reacts with DTNB through its thiol group to form 5-thionitrobenzoic acid. The reaction was monitored colorimetrically by measuring absorbance at 412 nm wavelength over 15 minutes. Enzyme activity was expressed as nmoles per minute per mg protein.

#### β-hydroxyacyl CoA dehydrogenase activity

β-hydroxyacyl CoA dehydrogenase (β-HAD) activity was measured in lysates prepared from frozen gastrocnemius muscles, as described previously^36^ with modifications. Briefly, gastrocnemius muscle lysate samples (standardized protein concentration to 7.5 mg/ml) were placed in triplicate wells in a 96-well plate containing 50 mM imidazole and 150 mM NADH as the assay medium. The reaction was initiated by adding acetoacetyl-CoA to a final concentration of 100 μM, and NADH disappearance was monitored spectrophotometrically by measuring the absorbance at 340 nm wavelength over a period of 7 min. Enzyme activity was expressed as nanomoles per minute per milligram of protein.

#### NAD^+^/NADH content assay

NAD^+^ and NADH levels in skeletal muscles were assessed in homogenate samples of frozen quadriceps muscles as previously described^37^. Briefly, Tissue lysates were subjected to an acid/base extraction procedure, followed by the measurement of NAD^+^ and NADH content through utilizing an enzyme cycling-based (alcohol dehydrogenase II) that is coupled to the reduction of methylthiazolyldiphenyl-tetrazolium bromide, which is colorimetrically monitoring at a wavelength of 570 nm. NAD^+^ and NADH content was expressed as nmol/g tissue.

#### Respiratory control ratio assay

Mitochondrial oxygen consumption was assessed in fresh saponin-permeabilized soleus muscle fibers using a Clark oxygen electrode connected to an Oxygraph Plus recorder (Hansatech Instruments Ltd., Norfolk, England) following the previously described methods^38,39^. The freshly excised soleus muscles from HSA^*Cre*^ and PDH^SkM-/-^ mice were immediately placed in an ice-cold isolation buffer (in mmol/L: 2.77 Ca-K2EGTA, 7.23 K2EGTA, 3 K2HPO4, 9.5 MgCl2, 5.7 Na2ATP, 15 phosphocreatine, 20 imidazole, 20 taurine, 0.5 dithiothreitol, and 49 K-methanesulfonate, 1 μM leupeptin, pH 7.1, at 0°C). The muscle was finely dissected into pieces and separated into fibers using forceps under a dissecting microscope in ice-cold isolation buffer. Subsequently, the fibers were permeabilized with 50 μg/mL saponin for 30-min at 4°C and washed three times for 5-min each in ice-cold respiration buffer (0.5 mM EGTA, 3 mM MgCl2.6H2O, 20 mM taurine, 10 mM KH2P04, 20 mM HEPES, 1 g liter-1 BSA, 60 mM potassium-lactobionate, 110 mM mannitol, 0.3 mM dithiothreitol). The fibers were then transferred to a respiration chamber containing 1.8 mL respiration buffer, and the rate of oxygen consumption was measured before and after adding 2 mM ADP with 5 mM malate and 10 mM glutamate as respiratory substrates to initiate basal respiration. The respiratory control ratio (RCR), which estimates mitochondrial respiration efficiency, was calculated as the ratio of basal to ADP-stimulated respiration rates.

### QUANTIFICATION AND STATISTICAL ANALYSIS

#### Statistical analysis

All values are presented as means ± standard error of the mean (SEM). Statistical significance was assessed using unpaired Student’s *t*-tests or repeated measures ANOVA followed by a Bonferroni *post-hoc* analysis, as appropriate, and as indicated in the figure legends. Differences were considered significant when *P* < 0.05. No data were considered outliers by testing or were arbitrarily excluded. Sample sizes were specified in the figure legends, and at least 5 mice per group were used for *in vivo* experiments. The operator(s) responsible for performing the metabolomic profiling were blinded to the genotype/treatment information. GraphPad Prism 9 software was used for all data analysis.

## DATA AND CODE AVAILABILITY

This study did not generate/analyze any datasets/code.

