## Supplementary figures and images for "Loss of Skeletal Muscle Pyruvate Dehydrogenase Induces Lactic Acidosis and Adaptive Anaplerotic Compensation via Pyruvate-Alanine Cycling and Glutaminolysis"

### Supplementary Figures

#### Supplementary Figure 1

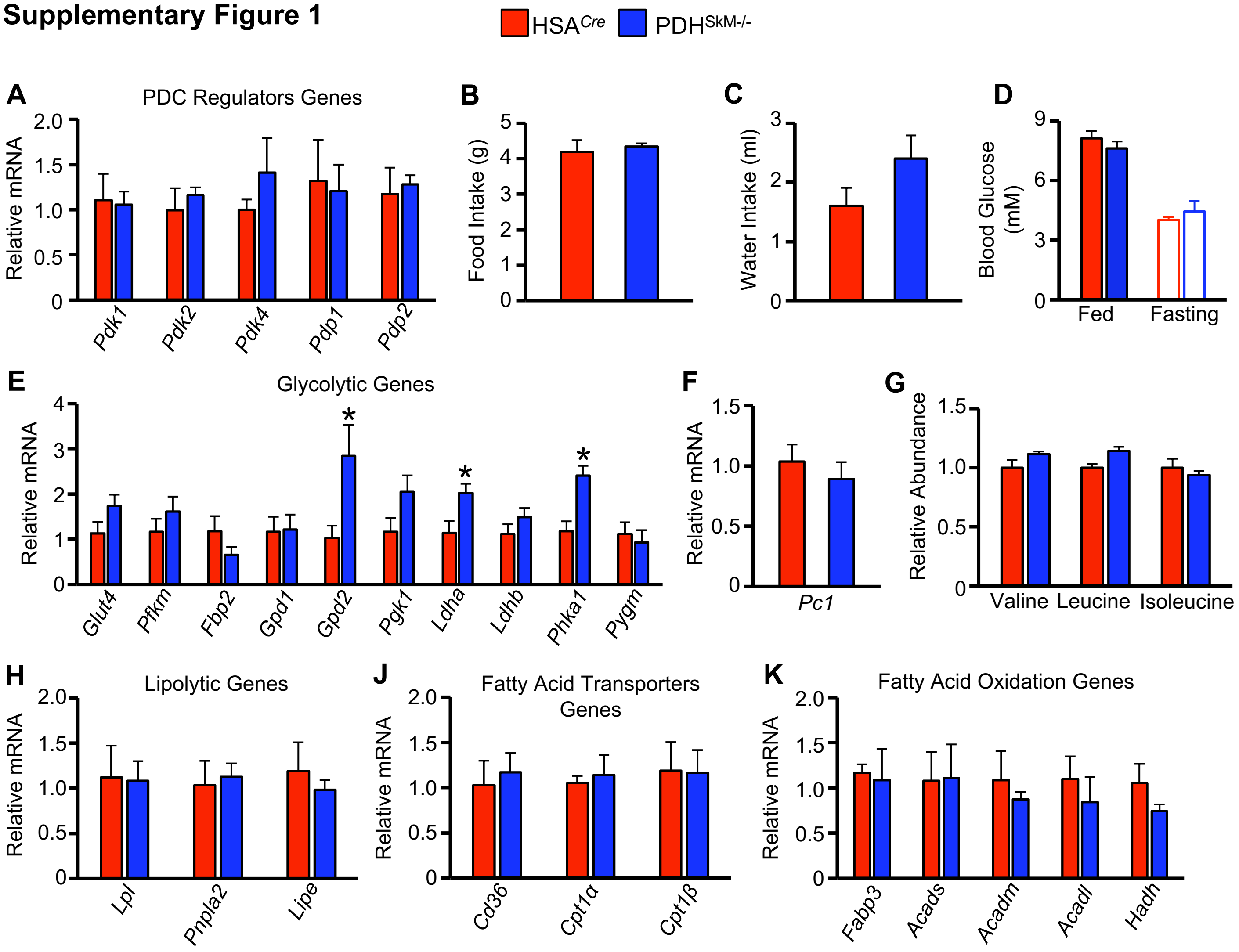

#### Supplementary Figure 2

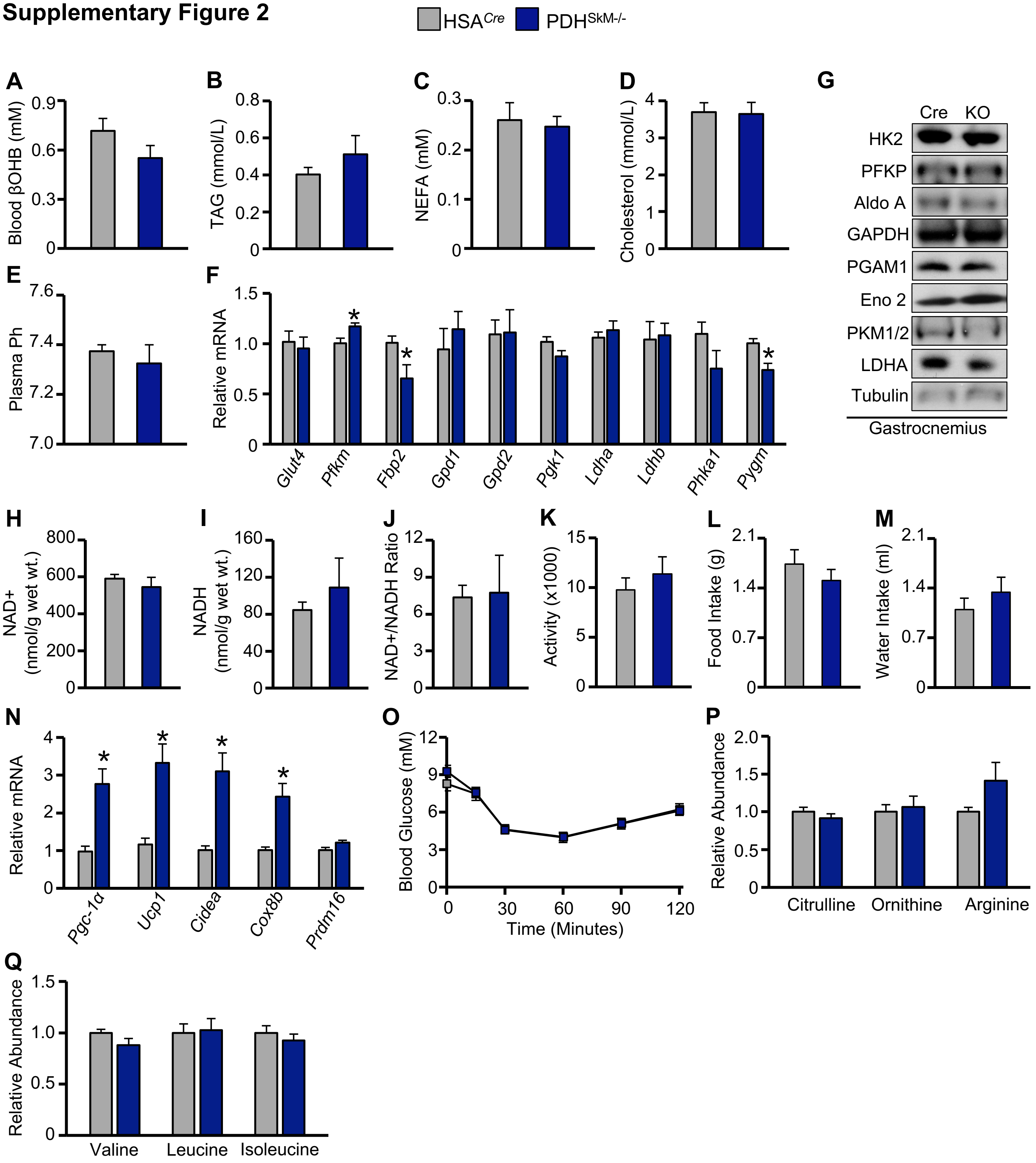
